# nf-core/mag: a best-practice pipeline for metagenome hybrid assembly and binning

**DOI:** 10.1101/2021.08.29.458094

**Authors:** Sabrina Krakau, Daniel Straub, Hadrien Gourlé, Gisela Gabernet, Sven Nahnsen

## Abstract

The analysis of shotgun metagenomic data provides valuable insights into microbial communities, while allowing resolution at individual genome level. In absence of complete reference genomes, this requires the reconstruction of metagenome assembled genomes (MAGs) from sequencing reads. We present the nf-core/mag pipeline for metagenome assembly, binning and taxonomic classification. It can optionally combine short and long reads to increase assembly continuity and utilize sample-wise group-information for co-assembly and genome binning. The pipeline is easy to install - all dependencies are provided within containers -, portable and reproducible. It is written in Nextflow and developed as part of the nf-core initiative for best-practice pipeline development. All code is hosted on GitHub under the nf-core organization https://github.com/nf-core/mag and released under the MIT license.

## INTRODUCTION

Shotgun metagenomic approaches enable genomic analyses of all microbes within, for example, environmental or host-associated microbiome communities. Since most bacteria cannot be cultured, isolated and individually sequenced, reference databases for microbial genomes are often incomplete. Thus, one of the main tasks in metagenomic data analysis is to reconstruct the individual genomes directly from the given mixture of metagenomic reads.

A typical reference-independent metagenomic workflow consists of preprocessing raw reads, assembly and binning to generate so-called metagenome assembled genomes (MAGs), as well as taxonomic and functional annotation. However, assemblies based on short reads often suffer from being highly fragmented. In contrast, long reads can be used to generate continuous assemblies, but suffer from high error rates. Hybrid assembly approaches combine the advantages of both short and long reads and thus, can produce continuous and accurate assemblies at reasonable cost (1). Another important aspect is whether to combine information across samples for assembly. When analysing multiple samples that contain the same microbes (e.g. cultures, time-series), co-assembly increases sequencing depth and can thus improve assembly completeness, in particular with respect to low abundant genomes. On the other hand, co-assembly can increase the metagenome complexity and result in more fragmentation or hybrid contigs of multiple similar genomes (2, 3).

Several pipelines have been developed for the assembly and binning of metagenomes (4, 5, 6). However, only a few pipelines such as Muffin (7) and ATLAS (8) make use of workflow management systems, such as Snakemake (9) or Nextflow (10), which facilitate scalability, portability, reproducibility and ease of application. These pipelines have different strengths and weaknesses, but only Muffin supports hybrid assembly and none of them supports co-assembly.

Here we introduce nf-core/mag, a pipeline for hybrid assembly of metagenomes, binning and taxonomic classification of MAGs. It allows the use of group information to perform co-assembly and the computation of co-abundances used for genome binning. The pipeline is written in Nextflow and part of the nf-core collection of community curated best-practice pipelines (11).

## MATERIAL AND METHODS

### Implementation and reproducibility

nf-core/mag is written in Nextflow, making use of the new DSL2 syntax (see Supplementary Material: Section 2). The pipeline strongly benefits from the nf-core framework, which enforces a set of strict best-practice guidelines to ensure high-quality pipeline development and maintenance (11). For example, pipelines must provide comprehensive documentation as well as community support via GitHub Issues and dedicated Slack channels. Pipeline portability and reproducibility are enabled through 1) pipeline versioning (i.e. tagged releases on GitHub), 2) building and archiving associated containers that contain the required software dependencies (i.e. the exact same compute environment can be used over time and across systems) and 3) a detailed reporting of the used pipeline/software versions and applied parameters. The use of container technologies such as Docker and Singularity enable reproducibility and portability also across different compute systems, i.e. local computers, HPC clusters and cloud platforms. The nf-core/mag pipeline comes with a small test dataset that is used for continuous integration (CI) testing with GitHub Actions. In addition, “full-size” pipeline tests are run on AWS for each pipeline release to ensure cloud compatibility and an error-free performance on real-world datasets. The full-size test results for each pipeline release are displayed on the nf-core website (https://nf-co.re/mag/results).

Although the nf-core framework facilitates reproducibility at several layers, it is further crucial to ensure that the individual tools that are part of the pipeline can be run in a deterministic and reproducible manner. For this purpose, nf-core/mag offers dedicated reproducibility settings, for example, to set a random seed parameter, to fix and report multi-threading parameters as well as to generate and/or save required databases, whose public versions do not always remain accessible (see Supplementary Material: Section 3).

### Simulation of metagenomic data

To show exemplary results generated with the nf-core/mag pipeline, we simulated metagenomic time series data with CAMISIM (12). For this, CAMISIM was applied based on the genome sources from the “CAMI II challenge toy mouse gut dataset” (13), containing 791 genomes, while set to generate Illumina and Nanopore reads. Two groups of samples were simulated, each comprising a time series of four samples (for details see Supplementary Material: Section 4). The simulated datasets as well as a sample sheet file, which can be used as input for the nf-core/mag pipeline, are available at https://doi.org/10.5281/zenodo.5155395.

## RESULTS

### Pipeline overview

An overview of the nf-core/mag pipeline is shown in Figure 1A. The input can be either directly provided FASTQ files containing the short reads or a sample sheet in CSV format containing the paths to short and, optionally, long read files as well as additional group information.

**Figure 1.**
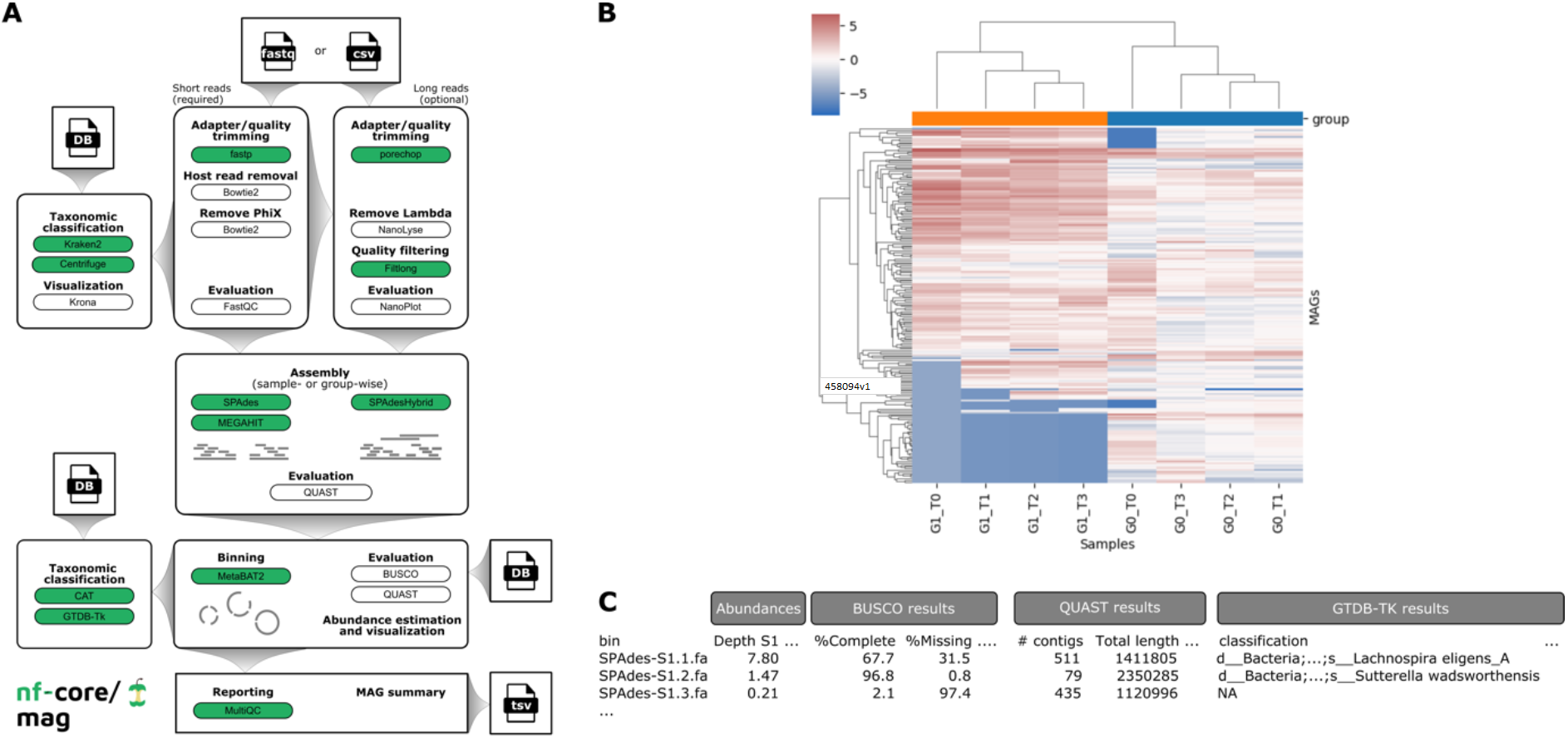
**A**) Overview of the nf-core/mag pipeline (v2.1.0). **B**) Clustered heatmap showing MAG abundances, i.e. centered log-ratio depths across samples. **C**) Schematic representation of MAG summary output, containing abundance information, QC metrics and taxonomic classifications.

#### Pre-processing

The pipeline starts with preprocessing the raw reads. For short Illumina reads, fastp (14) is used for adapter and quality trimming, Bowtie2 (15) for identifying and removing host or PhiX reads, and FastQC (https://www.bioinformatics.babraham.ac.uk/projects/fastqc/) for quality control (QC) on the raw and preprocessed reads. For long Nanopore reads, porechop (https://github.com/rrwick/Porechop) is used for adapter trimming, NanoLyse (https://github.com/rrwick/Filtlong) to remove phage lambda control contamination, and Filtlong (16) for quality filtering (host read contamination is indirectly removed based on the filtered short reads). QC on the raw and processed long reads is performed using NanoPlot (https://github.com/rrwick/Filtlong).

#### Assembly and binning

The preprocessed reads are then de novo assembled using MEGAHIT (17) or SPAdes (18). If both short and long reads are provided, a hybrid assembly can be performed using hybridSPAdes (19). By default, nf-core/mag assembles the reads of each sample individually. However, it provides the option to compute co-assemblies according to user specified group information. MetaBAT2 (20) is then used to bin the contigs into individual MAGs based on nucleotide frequencies and co-abundance patterns across samples (within the same group, by default). The pipeline further estimates MAG abundances for the different samples from contig sequencing depths. QUAST (21) summarizes QC features of the generated assemblies and MAGs. MAG completeness and contamination is estimated by BUSCO (22), which makes use of near-universal single-copy orthologs.

#### Taxonomic classification

Finally, MAGs are taxonomically annotated using GTDB-TK (23) or CAT/BAT (24). While CAT/BAT is able to taxonomically classify any MAG, GTDB-TK requires a number of marker genes and is therefore only applied to MAGs passing quality thresholds regarding the completeness and contamination priorly estimated with BUSCO. Besides the results from the individual tools, nf-core/mag outputs a summary containing estimated abundances, as well as the main QUAST, BUSCO and GTDB-TK metrics for each MAG (see Figure 1C).

#### Quality assurance

Preprocessed short reads are classified using Kraken2 (25) or Centrifuge (26) and visualized in Krona charts (27) to assess potential contamination and the microbial community before the assembly. MultiQC (28) is used to generate a comprehensive quality report aggregating the QC results across all samples.

For a comparison to existing pipelines for metagenome assembly and binning see Supplementary Table S1.

### Exemplary results

We ran nf-core/mag v2.1.0 on the metagenomic time series data simulated with CAMISIM. Figure 1B shows an example heatmap representing MAG abundances across samples, obtained with nf-core/mag performing hybrid, group-wise co-assembly. To illustrate the possible impact of the assembly setting, we compared the results for four different nf-core/mag settings, i.e. 1) short read only, sample-wise assembly, 2) hybrid, sample-wise assembly, 3) short read only, group-wise co-assembly and 4) hybrid, group-wise co-assembly. Figure 2 shows a comparison of the resulting assemblies with respect to commonly used assembly metrics. The results demonstrate that - for this particular time series data - both hybrid assembly as well as group-wise co-assembly increase the assembly’s size, its N50 value and the number of reconstructed MAGs, and thus likely the overall assembly completeness. For further details and results see Supplementary Material: Section 5.

**Figure 2.**
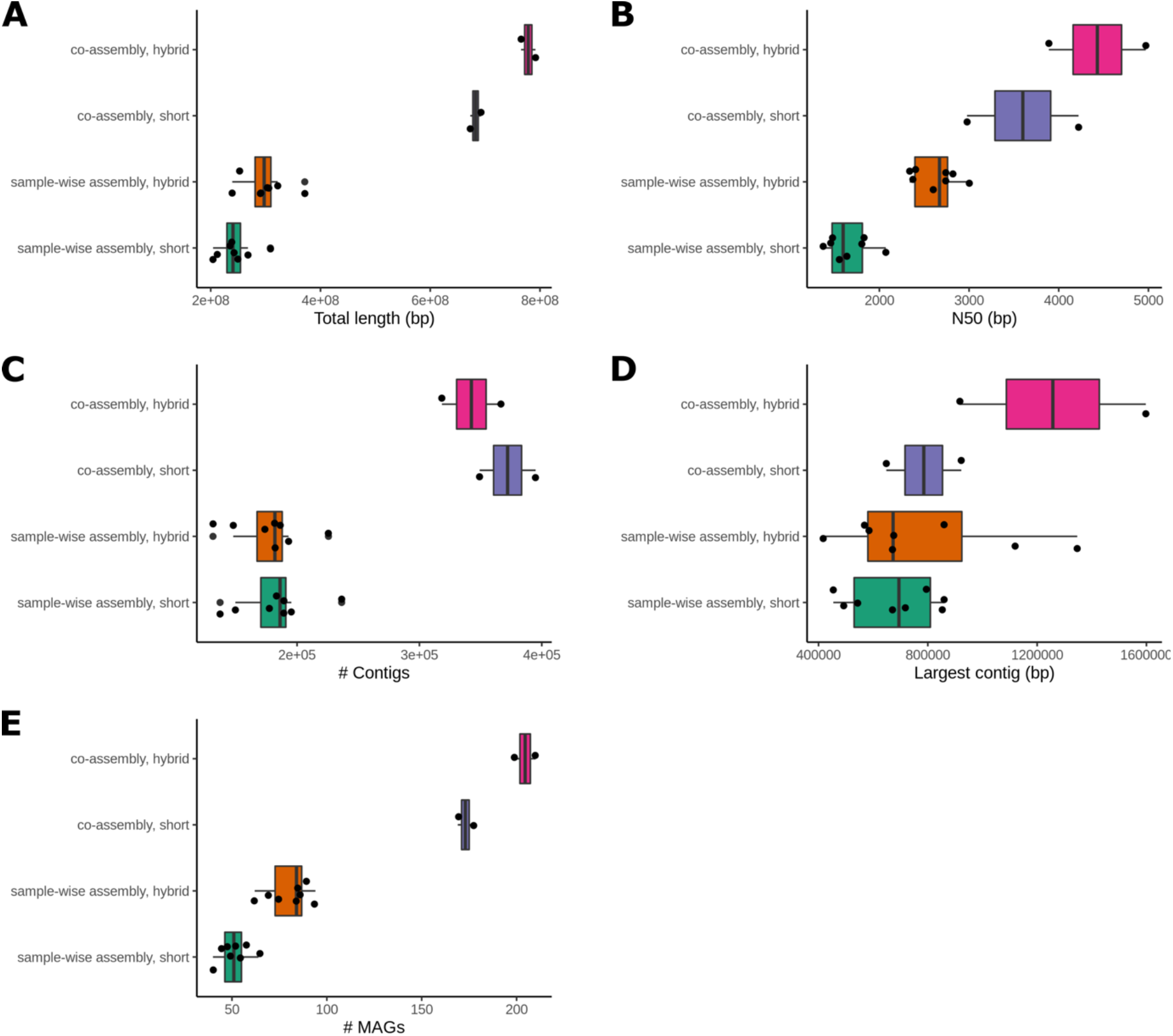
Assembly metrics obtained using different nf-core/mag assembly settings on the simulated data: sample-wise assembly, group-wise co-assembly, short read only assembly or hybrid assembly. Each point corresponds to one assembly, originating either from one sample or one group. Metrics displayed are **A**) total length of the assembly in base pairs, **B**) N50 value (i.e. the length of the shortest contig that needs to be included to cover least 50% of the genome), **C**) number of contigs in the final assembly, **D**) size of the largest contig in base pairs, and **E**) number of MAGs identified in the final assembly. The metrics **A**) – **D**) are part of the QUAST assembly summary.

## DISCUSSION

We implemented nf-core/mag, an easy to install, reproducible and portable pipeline for hybrid assembly, binning and taxonomic classification of metagenomes. It provides comprehensive usage documentation (https://nf-co.re/mag) and community support via a dedicated Slack channel (https://nfcore.slack.com/channels/mag). One important advantage over existing pipelines is that it can utilize sample-wise group information to perform co-assembly and/or to compute co-abundances used for the genome binning step. This enables users to choose the approach most suitable to their specific research question and experimental setup, or even to compare different settings.

The pipeline was already successfully applied in microbial studies (29, 30) and, as part of nf-core, will be constantly maintained and developed further to keep up with state-of-the-art analysis methods. Moreover, the modular DSL2 structure allows future integrations with other (sub-)workflows, for example, for the analysis of metatranscriptomic data.

## ACKNOWLEDGEMENT

We thank the nf-core community for general support during the writing of the pipeline.

## REFERENCES

1. Overholt, W. A., Hölzer, M., Geesink, P., Diezel, C., Marz, M., & Küsel, K. (2020). Inclusion of Oxford Nanopore long reads improves all microbial and viral metagenome-assembled genomes from a complex aquifer system. Environ. Microbiol., 22(9), 4000–4013.

2. Hofmeyr, S., Egan, R., Georganas, E., Copeland, A. C., Riley, R., Clum, A., Eloe-Fadrosh, E., Roux, S., Goltsman, E., Buluç, A., Rokhsar, D., Oliker, L., & Yelick, K. (2020). Terabase-scale metagenome coassembly with MetaHipMer. Sci. Rep., 10(1), 10689.

3. Olm, M. R., Brown, C. T., Brooks, B., & Banfield, J. F. (2017). dRep: A tool for fast and accurate genomic comparisons that enables improved genome recovery from metagenomes through de-replication. ISME J., 11(12), 2864–2868.

4. Fourquet, J., Noirot, C., Klopp, C. C., Pinton, P., Combes, S., Hoede, C., & Pascal, G. (2020). Whole metagenome analysis with metagWGS [Poster]. JOBIM2020.

5. Tamames, J., & Puente-Sánchez, F. (2019). SqueezeMeta, A Highly Portable, Fully Automatic Metagenomic Analysis Pipeline. Front. Microbiol., 0.

6. Uritskiy, G. V., DiRuggiero, J., & Taylor, J. (2018). MetaWRAP—a flexible pipeline for genome-resolved metagenomic data analysis. Microbiome, 6(1), 158.

7. Van Damme, R., Hölzer, M., Viehweger, A., Müller, B., Bongcam-Rudloff, E., & Brandt, C. (2021). Metagenomics workflow for hybrid assembly, differential coverage binning, metatranscriptomics and pathway analysis (MUFFIN). PLoS Comput. Biol., 17(2), e1008716.

8. Kieser, S., Brown, J., Zdobnov, E. M., Trajkovski, M., & McCue, L. A. (2020). ATLAS: A Snakemake workflow for assembly, annotation, and genomic binning of metagenome sequence data. BMC Bioinformatics, 21(1), 257.

9. Köster, J., & Rahmann, S. (2018). Snakemake—A scalable bioinformatics workflow engine. Bioinformatics, 34(20), 3600–3600.

10. Di Tommaso, P., Chatzou, M., Floden, E. W., Barja, P. P., Palumbo, E., & Notredame, C. (2017). Nextflow enables reproducible computational workflows. Nat. Biotechnol., 35(4), 316–319.

11. Ewels, P. A., Peltzer, A., Fillinger, S., Patel, H., Alneberg, J., Wilm, A., Garcia, M. U., Di Tommaso, P., & Nahnsen, S. (2020). The nf-core framework for community-curated bioinformatics pipelines. Nat. Biotechnol., 38(3), 276–278.

12. Fritz, A., Hofmann, P., Majda, S., Dahms, E., Dröge, J., Fiedler, J., Lesker, T. R., Belmann, P., DeMaere, M. Z., Darling, A. E., Sczyrba, A., Bremges, A., & McHardy, A. C. (2019). CAMISIM: Simulating metagenomes and microbial communities. Microbiome, 7(1), 17.

13. Meyer, F., Lesker, T.-R., Koslicki, D., Fritz, A., Gurevich, A., Darling, A. E., Sczyrba, A., Bremges, A., & McHardy, A. C. (2021). Tutorial: Assessing metagenomics software with the CAMI benchmarking toolkit. Nat. Protoc., 16(4), 1785–1801.

14. Chen, S., Zhou, Y., Chen, Y., & Gu, J. (2018). fastp: An ultra-fast all-in-one FASTQ preprocessor. Bioinformatics, 34(17), i884–i890.

15. Langmead, B., & Salzberg, S. L. (2012). Fast gapped-read alignment with Bowtie 2. Nat. Methods, 9(4), 357–359.

16. De Coster, W., D’Hert, S., Schultz, D. T., Cruts, M., & Van Broeckhoven, C. (2018). NanoPack: Visualizing and processing long-read sequencing data. Bioinformatics, 34(15), 2666–2669.

17. Li, D., Luo, R., Liu, C.-M., Leung, C.-M., Ting, H.-F., Sadakane, K., Yamashita, H., & Lam, T.-W. (2016). MEGAHIT v1.0: A fast and scalable metagenome assembler driven by advanced methodologies and community practices. Methods, 102, 3–11.

18. Nurk, S., Meleshko, D., Korobeynikov, A., & Pevzner, P. A. (2017). metaSPAdes: A new versatile metagenomic assembler. Genome Res., 27(5), 824–834.

19. Antipov, D., Korobeynikov, A., McLean, J. S., & Pevzner, P. A. (2016). hybridSPAdes: An algorithm for hybrid assembly of short and long reads. Bioinformatics, 32(7), 1009–1015.

20. Kang, D. D., Li, F., Kirton, E., Thomas, A., Egan, R., An, H., & Wang, Z. (2019). MetaBAT 2: An adaptive binning algorithm for robust and efficient genome reconstruction from metagenome assemblies. PeerJ, 7, e7359.

21. Gurevich, A., Saveliev, V., Vyahhi, N., & Tesler, G. (2013). QUAST: Quality assessment tool for genome assemblies. Bioinformatics, 29(8), 1072–1075.

22. Simão, F. A., Waterhouse, R. M., Ioannidis, P., Kriventseva, E. V., & Zdobnov, E. M. (2015). BUSCO: Assessing genome assembly and annotation completeness with single-copy orthologs. Bioinformatics, 31(19), 3210–3212.

23. Chaumeil, P.-A., Mussig, A. J., Hugenholtz, P., & Parks, D. H. (2020). GTDB-Tk: A toolkit to classify genomes with the Genome Taxonomy Database. Bioinformatics, 36(6), 1925–1927.

24. von Meijenfeldt, F. A. B., Arkhipova, K., Cambuy, D. D., Coutinho, F. H., & Dutilh, B. E. (2019). Robust taxonomic classification of uncharted microbial sequences and bins with CAT and BAT. Genome Biol., 20(1), 217.

25. Wood, D. E., Lu, J., & Langmead, B. (2019). Improved metagenomic analysis with Kraken 2. Genome Biol., 20(1), 257.

26. Kim, D., Song, L., Breitwieser, F. P., & Salzberg, S. L. (2016). Centrifuge: Rapid and sensitive classification of metagenomic sequences. Genome Res., 26(12), 1721–1729.

27. Ondov, B. D., Bergman, N. H., & Phillippy, A. M. (2011). Interactive metagenomic visualization in a Web browser. BMC Bioinformatics, 12(1), 385.

28. Ewels, P., Magnusson, M., Lundin, S., & Käller, M. (2016). MultiQC: Summarize analysis results for multiple tools and samples in a single report. Bioinformatics, 32(19), 3047–3048.

29. Huang, Y.-M., Straub, D., Blackwell, N., Kappler, A., & Kleindienst, S. (2021). Meta-omics Reveal Gallionellaceae and Rhodanobacter Species as Interdependent Key Players for Fe(II) Oxidation and Nitrate Reduction in the Autotrophic Enrichment Culture KS. Appl. Environ. Microbiol., 87(15), e00496–21.

30. Huang, Y.-M., Straub, D., Kappler, A., Smith, N., Blackwell, N., & Kleindienst, S. (2021). A Novel Enrichment Culture Highlights Core Features of Microbial Networks Contributing to Autotrophic Fe(II) Oxidation Coupled to Nitrate Reduction. Microb. Physiol., 1–16.

