## Supplementary Material for "nf-core/mag: a best-practice pipeline for metagenome hybrid assembly and binning"

Sabrina Krakau[1,*], Daniel Straub[1,2], Hadrien Gourlé[3], Gisela Gabernet[1] and Sven Nahnsen[1]

[1]Quantitative Biology Center (QBiC), University of Tübingen, Germany

[2]Microbial Ecology, Center for Applied Geosciences, University of Tübingen, Germany

[3]Department of Animal Breeding and Genetics, Swedish University of Agricultural Sciences, Uppsala, Sweden

#### 1. Comparison of existing pipelines for metagenome assembly and binning

Table S1 shows a comparison of nf-core/mag to existing pipelines that are implemented using workflow management systems, such as Snakemake or Nextflow, and allow a scalable and easy to use application on HPC clusters. Note that this comparison is focused on pipelines for the assembly and binning of metagenomes and does not include pipeline features for other analysis types.

| | **Feature** | **metagWGS v2.0** (1) | **Muffin v1.0.3** (2) | **ATLAS v2.6a2** (3) | **nf-core/mag v2.1.0** |
|---|---|---|---|---|---|
| | Workflow management system | Nextflow | Nextflow | Snakemake | Nextflow |
| Read QC | Supported sequencing technologies | Illumina | Illumina, Nanopore | Illumina | Illumina, Nanopore |
| | Adapter and quality Trimming | Cutadapt, Sickle | fastp, Filtlong | BBTools suite | fastp, Porechop, Filtlong |
| | PCR duplicate removal | BWA + Bedtools | No | BBTools suite | No |
| | Host-read removal | BWA + Bedtools | No | BBTools suite | Bowtie2 |
| Assembly | Short read assembly | metaSPAdes, MEGAHIT | metaSPAdes | metaSPAdes, MEGAHIT | metaSPAdes, MEGAHIT |
| | Long read assembly | No | MetaFlye | No | No |
| | Hybrid assembly | No | hybridSPAdes | hybridSPAdes (but no preprocessing for long reads) | hybridSPAdes |
| | Assembly refinement | Filters contigs based on mapped reads | MetaFlye assembly: Racon, Medeka, Pilon Reassembly after binning: Unicycler | Filters contigs based on mapped reads and contig lengths | No |

| | | | | | |
|---|---|---|---|---|---|
| | QC | metaQUAST | No | No | metaQUAST |
| | Group-wise co-assembly | No | No | No | Yes |
| Genome binning | Binning | MetaBAT2 | MetaBAT2, MaxBin2, CONCOCT | MetaBAT2, MaxBin2 | MetaBAT2 |
| | Group-wise co-abundances used for binning | No | No | Yes | Yes |
| | QC | BUSCO, metaQUAST (copied from nf-core/mag) | CheckM | CheckM | BUSCO, metaQUAST |
| | Bin refinement | No | MetaWRAP | DAS Tool | No |
| | Dereplication | No | No | dRep | No |
| | MAG abundance estimation | Yes | No | Yes | Yes |
| Annotation | Gene prediction | Prokka | No | Prodigal | Prodigal (run by BUSCO) |
| | Taxonomic classification | CAT/BAT | Sourmash with GTDB | GTDB-Tk | GTDB-Tk, CAT/BAT |
| | Functional annotation | eggNOG | eggNOG | eggNOG | No |
| Usability | Documentation | Yes | Yes | Yes (dedicated documentation of parameters missing) | Yes |
| | Aggregated QC results | MultiQC | No | No | MultiQC |
| | Interactive user support | GitLab Issues | GitHub Issues | GitHub Issues | GitHub Issues, Slack channel |
| | Reproducibility | No (no option to run MetaBAT2 in deterministic setting) | No (no option to run MetaBAT2 in deterministic setting) | No (no option to run MetaBAT2 in deterministic setting) | Yes |
| | Continuous integration tests | No | No | CircleCI | Small tests on GitHub Actions, "full-size" tests on AWS |
| | Launching and monitoring of pipeline runs via web interface | No | No | No | Nextflow tower (on local computers, cluster or cloud systems) |

**Table S1:** Comparison of metagenome assembly and binning pipelines.

### 2. nf-core and DSL2

All nf-core pipelines must be based on the nf-core template. This template was recently ported to the new Nextflow DSL2 syntax, which enables a modularised structure and reuse of components, with each process using its own BioContainer (4). nf-core/mag is ported to DSL2 since version 2.0.0. For a detailed description of the nf-core framework see the main nf-core publication (5) or the nf-core website (https://nf-co.re).

### 3. Reproducibility

Generating results that can be reproduced is a major challenge and many findings published in scientific literature can still not be replicated by other scientists (6). The nf-core framework enables reproducibility as described in the 'Material and Methods' section. To additionally ensure that the individual tools generate reproducible results, several reproducibility settings were implemented for nf-core/mag. MEGAHIT and SPAdes, for example, depend on multi-threading parameters and the number of CPUs used for computation can affect the final results. In nf-core/mag, the number of used CPUs can be fixed and reported accordingly to generate reproducible assemblies. This ensures that the specified number of CPUs is not increased in case these processes will be re-submitted (as is usually the case for nf-core pipelines, if the specified resource requirements for a process do not suffice). For MetaBAT2, a deterministic behaviour is enabled by default within this pipeline via a fixed seed parameter. Moreover, specific settings allow the generation and/or saving of databases for BUSCO or CAT, for which the required public databases do not always remain accessible.

### 4. Simulated metagenomic data

The metagenomic data was simulated with the most recent development version of CAMISIM at the time of preparing this article (available at https://doi.org/10.5281/zenodo.5137751)(7). Two groups of samples were generated by using different CAMISIM seeds ('seed=1000' and 'seed=1001'), each comprising a time series of four samples. To simulate Illumina reads, the parameters 'ncbi_taxdump=tools/ncbi-taxonomy_20180226.tar.gz', 'number_of_samples=4', 'genomes_total=500', 'genomes_real=500', 'mode=timeseries_lognormal', 'gauss_mu=1' and 'gauss_sigma=1' were specified in addition to the seed and the default parameters in the CAMISIM configuration file 'defaults/default_config.ini'. The simulated FASTQ datasets were split with respect to paired-end reads. For the simulation of Nanopore reads, CAMISIM was run in combination with NanoSim v2.5.0. The parameters 'anonymous=False', 'ncbi_taxdump=tools/ncbi-taxonomy_20180226.tar.gz', 'number_of_samples=4', 'genomes_total=500', 'genomes_real=500', 'mode=timeseries_lognormal', 'gauss_mu=1' and 'gauss_sigma=1' were specified in addition to the seed and the default parameters in the configuration file 'defaults/nanosim_config.ini'. For each sample the resulting genome-wise FASTQ files were merged.

### 5. Results on simulated data

We ran nf-core/mag with the different assembly settings on the simulated metagenomic data. The following command was used (Nextflow v21.04.1) to generate short read and hybrid sample-wise assemblies:

```
> nextflow run nf-core/mag -r 2.1.0 -profile cfc -c custom.config --input samplesheet.CAMISIM_hybrid.csv --
binning_map_mode all --spades_fix_cpus 40
--spadeshybrid_fix_cpus 40 --skip_megahit
```

Corresponding group-wise co-assemblies were generated with:

```
> nextflow run nf-core/mag -r 2.1.0 -profile cfc -c custom.config --input samplesheet.CAMISIM_hybrid.csv --
coassemble_group --binning_map_mode all
--spades_fix_cpus 40 --spadeshybrid_fix_cpus 40 --skip_megahit
```

A ‘custom.config’ file was used to increase the memory for SPAdes and contained:

```
process {
  withName: SPADES {
    memory      = 150.GB
  }
  withName: SPADESHYBRID {
    memory      = 150.GB
  }
}
```

Besides comparing the resulting assemblies (see Figure 2), we compared the reconstructed genomes with respect to commonly used MAG metrics. The results shown in Figure S1 demonstrate that the average number of contigs per MAG decreased with hybrid vs. short read assembly. However, the average quality across all MAGs does not increase when using the hybrid or co-assembly assembly setting. For example, performing group-wise co-assemblies results in a higher average contamination compared to sample-wise assemblies (see Figure S2 d)), and hybrid assemblies result in lower completenesses compared to short read assemblies (see Figure S2 c)). This is likely caused by the highly increased number of reconstructed MAGs when using these settings (see Figure 2E).

**a)**

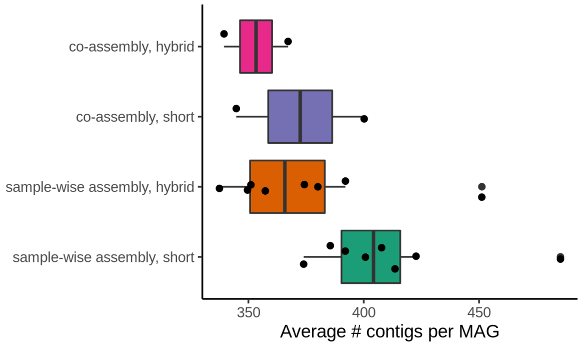


**b)**

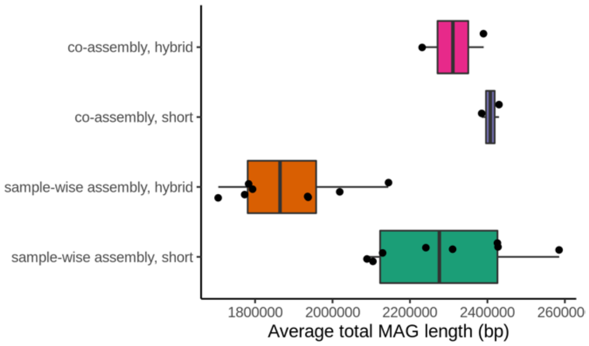

**c)** **d)**

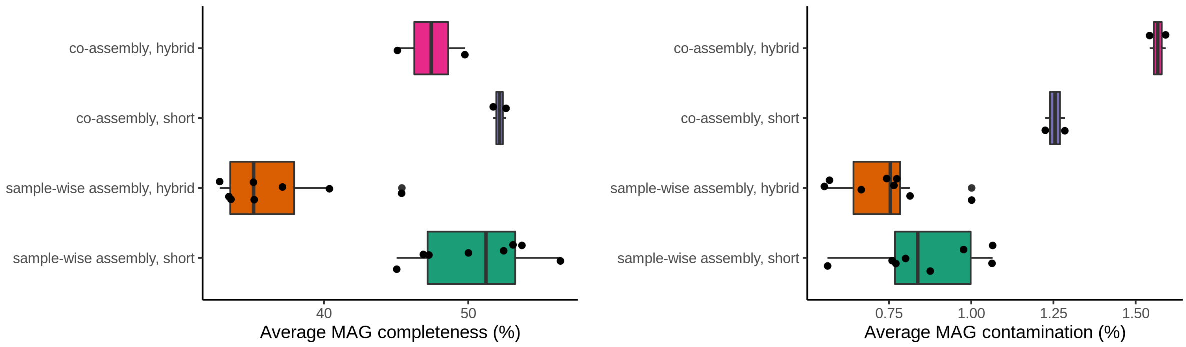


**Figure S1:** MAG-wise metrics obtained using different nf-core/mag assembly settings on the simulated data: sample-wise assembly, group-wise co-assembly, short read only assembly or hybrid assembly. Each point corresponds to one assembly, originating either from one sample or one group. Metrics displayed are **a)** average number of contigs per MAG, **b)** average total MAG length in base pairs, **c)** average MAG completeness and **d)** average MAG contamination. The number of contigs and the total length of each MAG were summarised by QUAST, the MAG completeness and contamination were estimated by BUSCO.

### References

1. Fourquet, J., Noirot, C., Klopp, C. C., Pinton, P., Combes, S., Hoede, C., & Pascal, G. (2020). Whole metagenome analysis with metagWGS [Poster]. *JOBIM2020*.
2. Van Damme, R., Hölzer, M., Viehweger, A., Müller, B., Bongcam-Rudloff, E., & Brandt, C. (2021). Metagenomics workflow for hybrid assembly, differential coverage binning, metatranscriptomics and pathway analysis (MUFFIN). *PLOS Computational Biology*, *17*(2), e1008716.
3. Kieser, S., Brown, J., Zdobnov, E. M., Trajkovski, M., & McCue, L. A. (2020). ATLAS: A Snakemake workflow for assembly, annotation, and genomic binning of metagenome sequence data. *BMC Bioinformatics*, *21*(1), 257.
4. da Veiga Leprevost, F., Grüning, B. A., Alves Aflitos, S., Röst, H. L., Uszkoreit, J., Barsnes, H., Vaudel, M., Moreno, P., Gatto, L., Weber, J., Bai, M., Jimenez, R. C., Sachsenberg, T., Pfeuffer, J., Vera Alvarez, R., Griss, J., Nesvizhskii, A. I., & Perez-Riverol, Y. (2017). BioContainers: An open-source and community-driven framework for software standardization. *Bioinformatics*, *33*(16), 2580–2582.
5. Ewels, P. A., Peltzer, A., Fillinger, S., Patel, H., Alneberg, J., Wilm, A., Garcia, M. U., Di Tommaso, P., & Nahnsen, S. (2020). The nf-core framework for community-curated bioinformatics pipelines. *Nature Biotechnology*, *38*(3), 276–278
6. Baker, M. (2016). 1,500 scientists lift the lid on reproducibility. *Nature*, *533*(7604), 452–454. https://doi.org/10.1038/533452a
7. Fritz, A., Hofmann, P., Belmann, P., Bremges, A., McHardy, A., Dröge, J., & DeMaere, M. (2021). skrakau/CAMISIM: Simulation of hybrid, time series data. *Zenodo*.
